# PIRATE: Plundering AlphaFold Predictions to Automate Protein Engineering

**DOI:** 10.1101/2024.10.02.611680

**Authors:** Graham Richardson, Jola Kopec, Carmen Esposito, Tim James

## Abstract

An important characteristic for many proteins is the presence of flexible loops or linkers known as intrinsically disordered regions. One downside to many of the state-of-the-art disorder prediction approaches is that they require significant computational time for each inference. Here, we introduce three novel surrogate models trained on AlphaFold2 predictions that rapidly encode local, regional, and global structural properties directly from primary sequence. We combined the outputs from these surrogate models, in an approach we term PIRATE, and show that this approach approximates the performance of AlphaFold2 for disorder prediction. Additionally, we show that PIRATE is much more sensitive to the effects of point mutants on disorder at distal sites than many current disorder prediction methods. Furthermore, we show that in the context of a greedy exploration algorithm, PIRATE’s ability to evaluate differences between point mutants makes it ideal for automating disorder-related protein engineering tasks.

## Introduction

A necessary requirement for many naturally occurring proteins is the ability to form interconverting conformational states where each of these conformational states is more entropically stable than intermediate forms. This characteristic tends to be an emergent requirement for molecular machines as organism complexity increases (Tompa et al., 2006) and it requires an inherent level of flexibility within specific segments of the molecule. In the context of proteins, these segments are termed intrinsically disordered regions (IDRs) (Tompa, 2002). Encoded in these IDRs is the tendency to form a wide range of rapidly fluctuating structures known as ensembles rather than stable three-dimensional structures (Lotthammer et al., 2024). This provides flexibility for adjacent segments to move or change conformation in precisely regulated ways (Vymětal et al., 2019). IDRs are also known to facilitate protein binding site function and appear to regulate higher order complex formation (Seoane & Carbone, 2021).

In 2021, the first Critical Assessment of protein Intrinsic Disorder prediction (CAID) experiment was published (Necci et al., 2021). Forty-three disorder prediction methods were validated across several disorder prediction tasks. The following year, researchers from the same group demonstrated that the protein structure prediction model AlphaFold2 (AF2) could be used to identify disordered regions in proteins using the model’s predicted local distance difference test (pLDDT) and inferred relative solvent accessibility (RSA) for protein residues (Piovesan et al., 2022). For the task of disorder prediction, with the original CAID dataset AF2 proved to be competitive with state-of-the-art approaches in the field. When the results of the CAID2 experiment were published, they included an evaluation of the AF2-RSA and AF2-pLDDT methodologies (Conte et al., 2023). Again, AF2 predictions proved to be top performers at several disorder prediction tasks. However, one clear drawback to using AF2 for disorder prediction is the considerable time and computational resources required to make each inference.

To create a more efficient disorder prediction method, we implemented a segmentation modeling approach, where the models were trained on AF2 predictions. The ability to accurately classify discrete segments of images (also known as semantic segmentation) is an underlying component to many modern computer vision artificial intelligence approaches (Chai et al., 2021). In the field of semantic segmentation, the first U-Net model represented a significant improvement over contemporary sliding-window based convolutional approaches (Ronneberger et al., 2015). The U-Net architecture is comprised of nearly symmetrical contracting and expanding pathways forming an hourglass shape.

However, the true innovation of U-Net was the creation of semantic pathways known as “skip connections” between corresponding layers of the contracting and expanding pathways. These skip connections allow the model to make random elastic deformations to the information within the contracting/expanding pathways while retaining feature maps of the information within the semantic pathways of the network. This property of U-Net allows the network to efficiently model nuances of training data with a minimal number of examples. Recent innovations to the U-Net model have focused on constructing denser (Zhou et al., 2018) or more efficient (Huang et al., 2020) semantic pathways within the network.

In this paper, we introduce three novel surrogate models that efficiently model local, regional, and global structural predictions from AF2. We combine these predictions in an approach we call Protein IDR Reduction Artificial inTElligence (PIRATE), which demonstrates comparable performance to AF2 at disorder prediction, but without the need for a GPU. PIRATE’s ability to infer how minor changes to a sequence will impact distal regions of disorder within that sequence makes it optimal for protein engineering tasks aimed at disorder reduction.

## Results

### Training AFSM (AlphaFold Surrogate Model) 1 with AF2 MAE and AFSM2 with AF2 pLDDT

Because AF2-predicted pLDDT or inferred RSA are sufficient to predict IDR’s, we hypothesized that creating surrogate models for AF2 outputs could prove to be a particularly powerful approach for efficient IDR classification. We collected AF2 predictions for 20 proteomes (Figure 1a) comprising 475,249 unique sequences from the AlphaFold API. Two outputs from AF2 predictions seemed immediately useful for predicting disorder: predicted alignment error and pLDDT. Predicted alignment error represents a global prediction of structural uncertainty, whereas pLDDT is the generated score of local structural confidence. We reasoned that having both a global and a local prediction of structural certainty would be practical for modeling the effects of small perturbations across large protein sequences.

**Figure 1.**
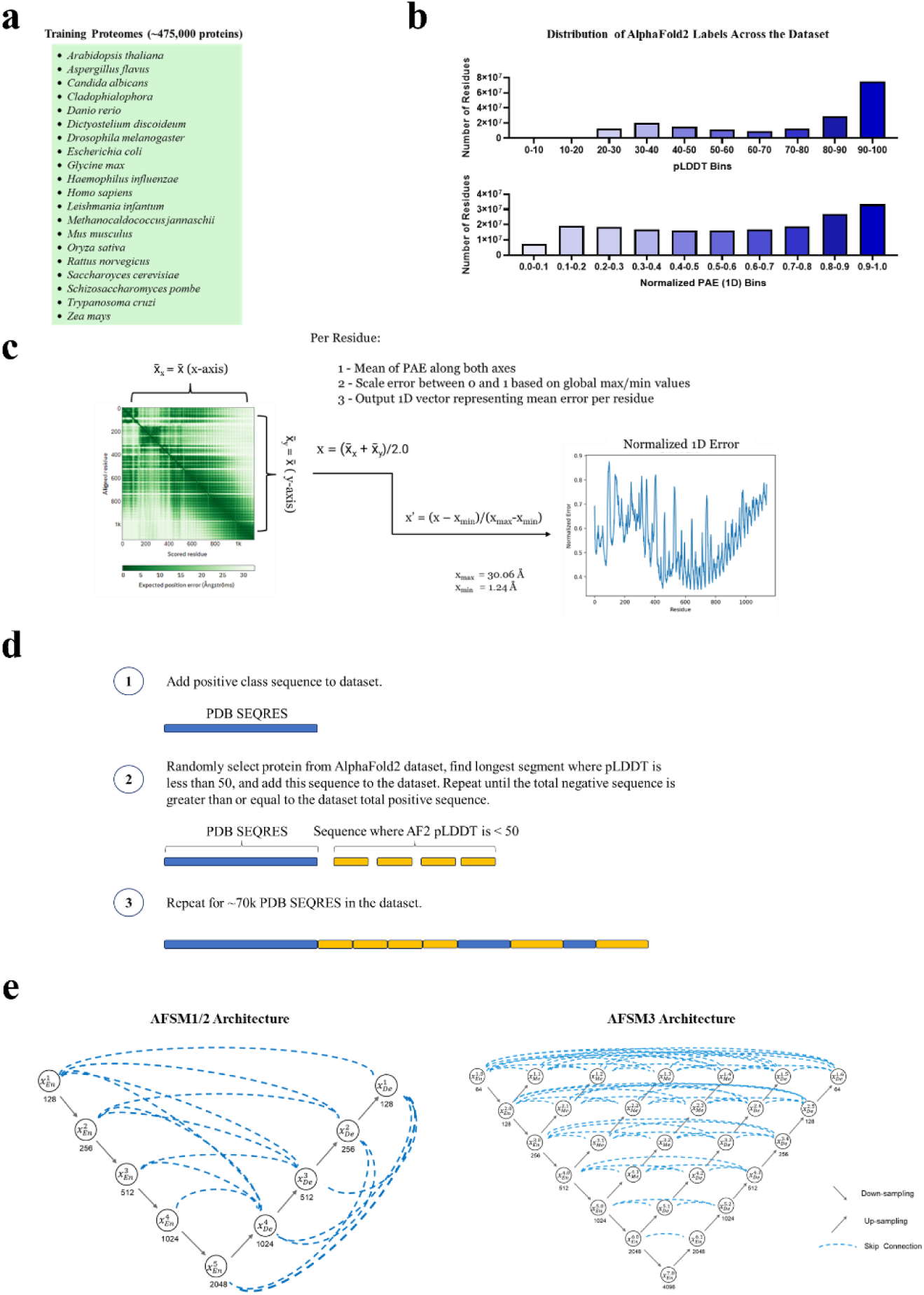
Creation and design of AF2 surrogate models. **(a)** A list of collected proteomes that comprise the AF2 dataset. **(b)** The distribution of pLDDT and one-dimensional alignment error across the dataset. **(c)** The method of converting AF2 predicted alignment error from a two-dimensional array to a one-dimensional vector. **(d)** Generation of a balanced dataset consisting of consisting of protein sequence from crystal structures and sequence that AF2 predicts to have a pLDDT < 0.5. **(e)** Diagram of the UNet+++ architecture used to create AFSM1 and AFSM2 and the UNet++ architecture used to create AFSM3.

For each sequence and each residue in every sequence, we compiled the predicted alignment error and pLDDT scores. AF2 predicted alignment error is a two-dimensional array representing the predicted distance uncertainty in angstroms for each residue in the sequence relative to every other residue. We converted these arrays into one-dimensional vectors by taking the mean error across each axis and then averaging over each residue (Figure 1c). The error was scaled using minimum and maximum values of the dataset. Within the dataset, we observed a general skew towards higher one-dimensional mean alignment error (MAE) and pLDDT scores on a per-residue basis (Figure 1b).

We tested multiple semantic segmentation approaches for modeling AF2 pLDDT and MAE outputs on a per-residue basis, and the U-Net3+ model described in Huang et al., 2020 demonstrated the lowest validation and testing error. Hereinafter, we will refer to the U-Net3+ trained models for MAE and pLDDT as AlphaFold Surrogate Models AFSM1 and AFSM2, respectively. The inventors of U-Net3+ hypothesized that the full-scale skip connections were particularly effective at retaining aggregated feature maps at different scales and we hypothesize this might cause U-Net3+ to be especially well-suited for segmenting disorder associated with widely varying sizes and distributions across protein domains (Figure 1e, AFMS1/2).

### Training AFSM3 with PDB SEQRES and AF2-predicted disordered sequences

While AFSM1 and AFM2 model global and local disorder using AF2-predicted alignment error and pLDDT, respectively, AFMS3 was designed to predict regional disorder, defined by the lack of a stable structure within specific protein segments. To train a model capable of learning regional positional confidence on a per-residue basis, we collected 70,549 protein sequences from PDB SEQRES records. These sequences, often being fragments of the full wild-type protein that a scientist was able to crystallize, served as positive examples of regional order. Although crystallographic structures sometimes include unresolved residues or regions, the PDB does not inherently contain negative examples, i.e., sequences that are unlikely to crystallize due to structural disorder. To construct these negative examples, we sampled from the previously described AF2 dataset (Figure 1a), selecting from a randomly chosen protein the longest continuous segment where each residue had a pLDDT lower than 50. This process of randomly selecting proteins and extracting disordered segments was repeated iteratively until the total amount of these negatively labeled residues was equal to or greater than the positively labeled residues in the dataset (Figure 1d), thus resulting in a balanced dataset.

This balanced classification dataset was used to train and test several semantic segmentation models for the classification of every residue in a protein sequence as either ordered or disordered. Among these, the U-Net++ model (Figure 1e, AFSM3) demonstrated the best performance and was therefore chosen for subsequent developments. Due to its dense, nested structure of skip layers, we hypothesize that U-Net++ is especially well-suited for modeling the highly variable intermediate range between local and global because of the minimized semantic gaps between the encoder and decoder pathways.

### PIRATE: A tool for efficient disorder prediction

The main outcome of this work is PIRATE (Protein IDR Reduction Artificial inTEllegence), a supervised classification model for predicting per-residue disorder in protein sequences. PIRATE encodes protein sequences using the predictions from the AFSM models described above. Unlike other existing disorder prediction methods, which perform well at predicting disorder observed in NMR experiments or identifying unmodeled residues in PDB structures (Necci et al., 2021), our model was specifically developed for detecting intrinsically disordered regions, i.e., regions that lack a stable three-dimensional structure. To accomplish this, we chose to train PIRATE using PDB data, where disordered sequences are underrepresented.

The training dataset was assembled by downloading X-ray crystallography PDB structures of single chain proteins. The residues from each PDB SEQRES were labeled as either modeled or unmodeled (disordered) according to whether they were present in the PDB structure. We predicted MAE, pLDDT, and regional probability of crystallization for each residue in these sequences using all three surrogate models. In addition, we encoded the SEQRES residues as ordinal values (Figure 2a).

**Figure 2.**
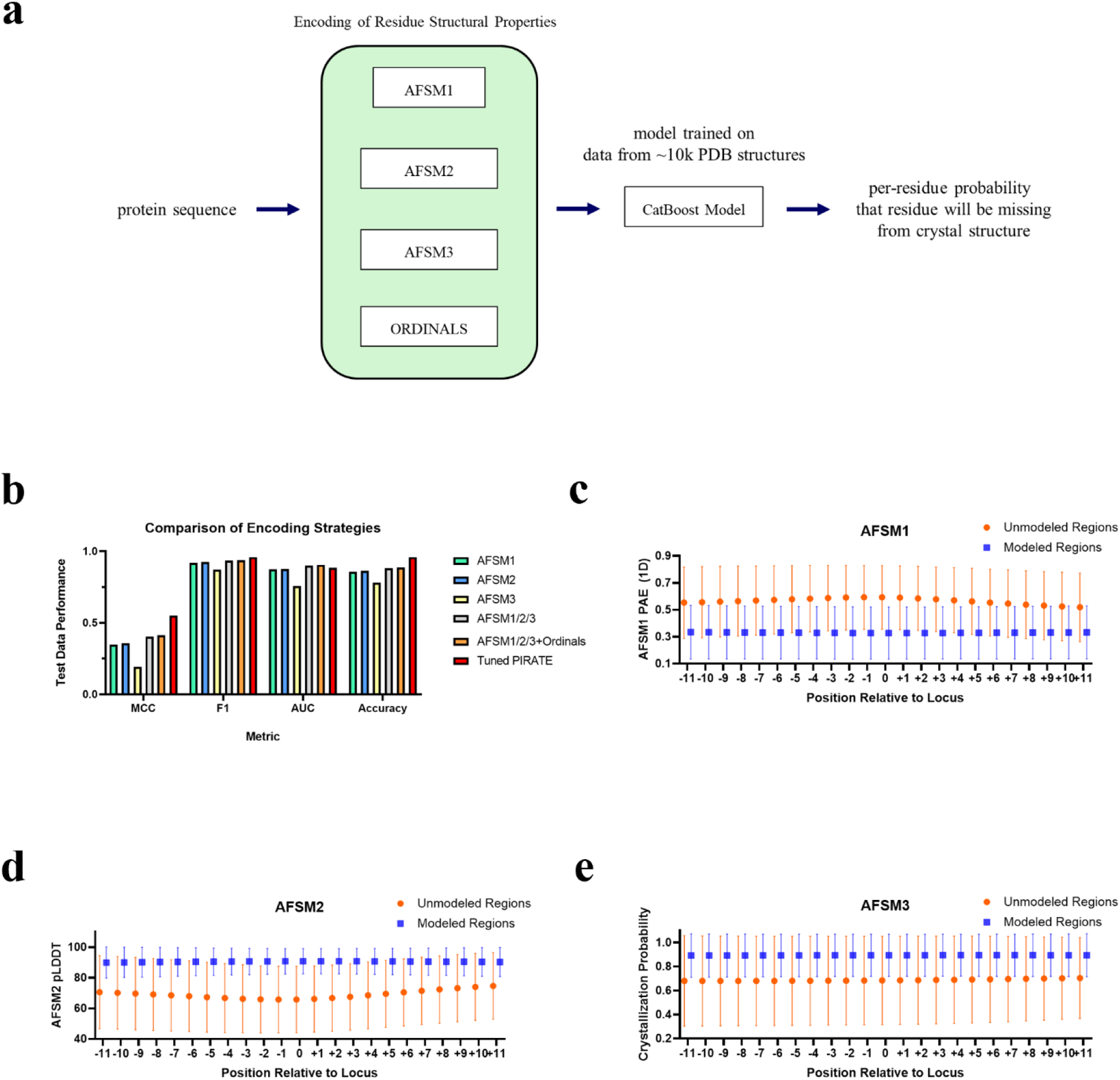
Creation and evaluation of PIRATE. **(a)** Diagram showing the design for PIRATE. **(b)** Performance of the Cat Boost model at predicting whether a residue is modeled or unmodeled using various encoding strategies. Error bars represent one standard deviation. **(c)** Plot showing a mean representation for outputs from AFSM1 for regions not modeled in the PDB dataset vs. those modeled in the PDB dataset. **(d)** Plot showing a mean representation for outputs from AFSM2 for regions not modeled in the PDB dataset vs. those modeled in the PDB dataset. **(e)** Plot showing a mean representation for outputs from AFSM3 for regions not modeled in the PDB dataset vs. those modeled in the PDB dataset.

These residue encodings were fed to a supervised learning model using a sliding window approach. For each residue, encodings from eleven residues on either side (total window size of 23 residues) were concatenated and used as input to a CatBoost (Dorogush et al., 2018) classifier (Figure 2a). We compared the performance of classifiers trained on individual ASFM model representations against those trained on combined encodings. The best performing model, which used a concatenation of all ASFM prediction encodings as well as ordinal values, achieved accuracy and F1 scores of 0.96 on a per-residue basis (Figure 2e). Building on these results, the final PIRATE model (Figure 2b) was established by optimizing the hyperparameters of this model with the Optuna library (Akiba et al., 2019).

To gain insight into how individual encodings contributed to the model’s predictions, we analyzed the predictions of each AFSM model for 23-residue windows centered on target residues, either modeled or unmodeled in the PDB data. When analyzing the mean predicted MAE from AFSM1 for windows of target unmodeled residues from the PDB dataset, we observed a peak MAE at the unmodeled residue that decreases gradually as residues become more distal (Figure 2c). Overall, the MAE of residue windows centered on unmodeled residues is much lower than that of windows centered on modeled residues. The opposite trend emerged from the AFSM2 predicted pLDDT for unmodeled windows: pLDDT is lowest at the target unmodeled residue and increases as distance from it grows (Figure 2d). As expected, the mean predicted pLDDT is higher across windows of modeled target residues.

The regional surrogate model, AFSM3, predicted lower crystallization probabilities for unmodeled residue windows compared to modeled ones. However, unlike the global (AFSM1) and local (AFSM2) models, AFSM3 did not exhibit a clear probabilistic change radiating outward from the target residue (Figure 2e). These trends in the surrogate model predictions serve as an important validation that the output of the surrogate models is aligned with the task of disorder prediction in our PDB dataset.

To further support these findings, we analyzed the residue prevalence within windows of both modeled and unmodeled target residues. As previously reported (Djinovic-Carugo & Carugo, 2015), we found a higher frequency of serine and glycine residues in disordered regions of PDB structures (Supplementary Figure 2).

### PIRATE performance evaluation on the CAID2 Disorder dataset

To assess the performance of PIRATE on a well-characterized dataset, we selected the recently published CAID2 Disorder-PDB dataset, where AlphaFold2 predictions achieved a second-place ranking (Conte et al., 2023). We validated that none of these sequences were present in the PDB dataset used for training our approach. PIRATE’s performance ranked third by two of the competition’s metrics (F1-max and APS) and fourth by AUC (Figure 3). Notably, while PIRATE performed slightly worse than the AlphaFold2-rsa method, it outperformed the AlphaFold2-pLDDT method (Piovesan et al., 2022). Additionally, while Spot-Disorder2 (Hanson et. al 2019) and AlphaFold-rsa (Piovesan et al., 2022) consistently performed better than PIRATE, inference with PIRATE takes between one and two seconds per sequence on a laptop CPU making it suitable for iterative sequence design/optimization tasks. Spot-Disorder2 and AlphaFold2 both have inference times more than 3 orders of magnitude greater than PIRATE, which we expect would make these approaches intractable for many protein design use cases.

**Figure 3.**
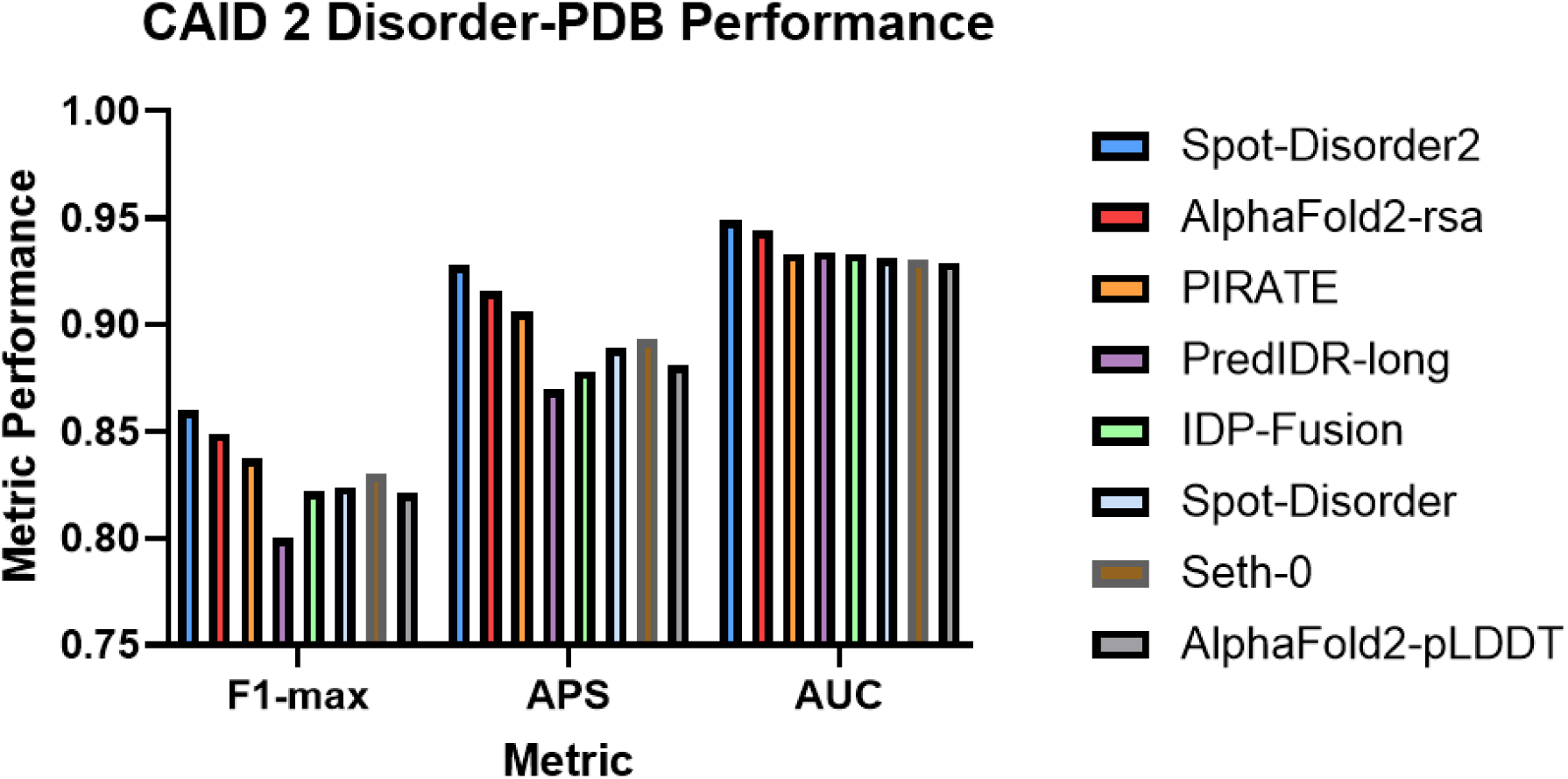
Evaluation of Disorder Prediction. **(a)** PIRATE performance on CAID2 Disorder-PDB dataset as compared to other top performing models from the competition.

### PIRATE predicts the effects of distal mutations on disorder

We speculated that to apply disorder prediction for protein design tasks, we would need an approach that is not only computationally efficient, but also able to predict small regions of instability with high specificity. One common approach for experimentally identifying such regions is partial trypsin digestion analysis. To see if PIRATE could replicate these experimental results, we performed PIRATE disorder prediction on the human proteins AADC and EF1a with experimentally characterized trypsin digestion sites (Bisello et al. 2023, Kinzy & Merrick, 1991) (Figure 4). We compared PIRATE’s results with two disorder prediction methods that have recently been utilized for protein design: DR-BERT (Alamdari et al, 2023) and IUPRED3 (Ferruz et al., 2022). Additionally, we benchmarked PIRATE’s predictions against some of the top-performing disorder prediction methods from the CAID2 competition: Spot-Disorder2 (Hanson et al., 2019), PredIDR-long (Han et al., 2024), IDP-Fusion (Tang et al., 2023), AF2-pLDDT (Piovesan et al., 2022), Seth-0 (Ilzhöfer et al., 2022), and Spot-Disorder (Hanson et al., 2017).

**Figure 4.**
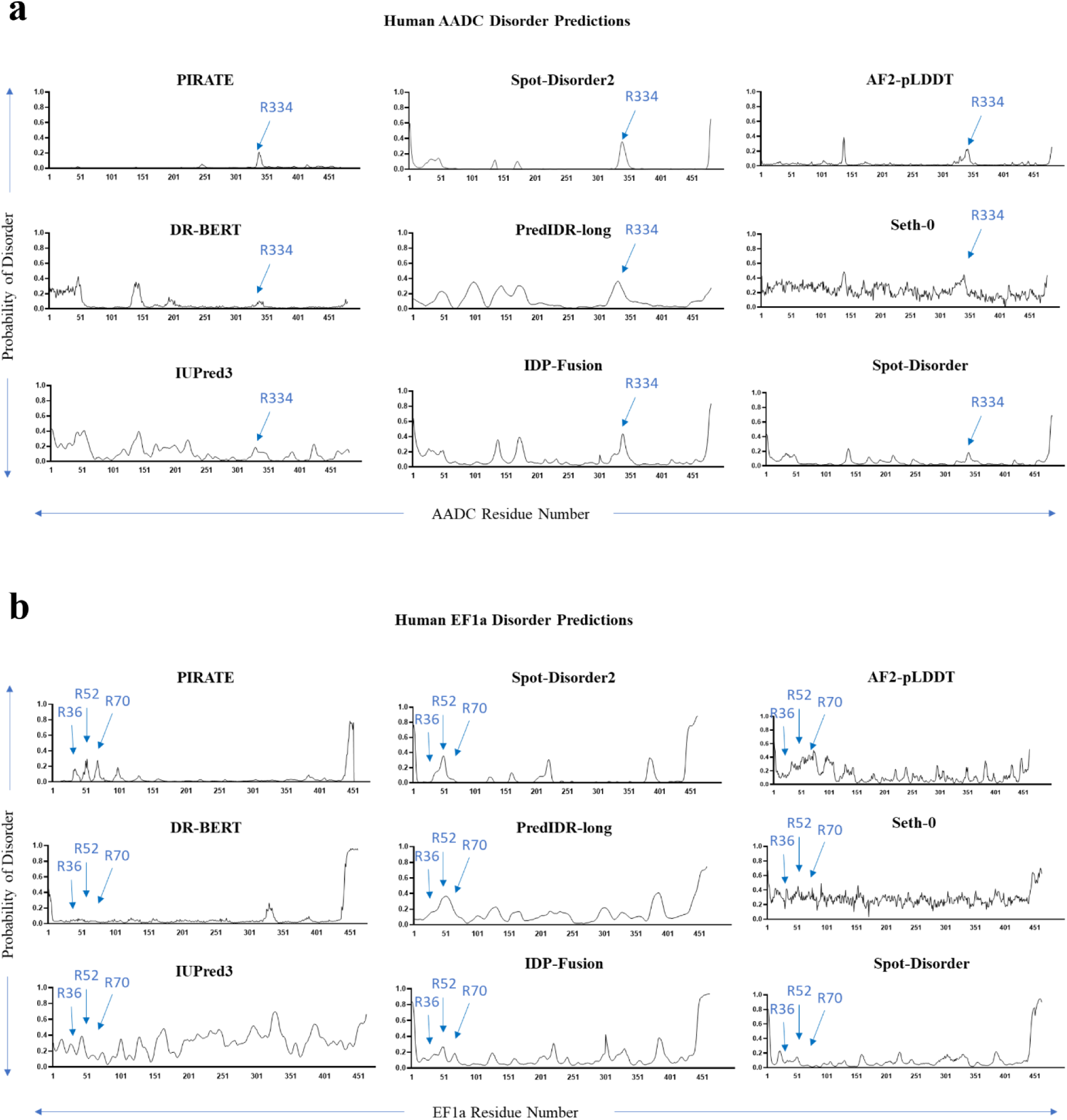
PIRATE Disorder Predictions Correlate with Trypsin Digestion Sites. **(a)** PIRATE demonstrates the ability to resolve the human AADC experimentally validated trypsin digestion site at R334 as a distinct peak of increased disorder probability. We compare this with disorder prediction methods previously used for protein design or methods that performed well in the CAID2 competition. **(b)** PIRATE demonstrates the ability to resolve the human EF1a experimentally validated trypsin digestion sites at R36, R52, and R70 as distinct peaks of increased disorder probability. We compare this with disorder prediction methods previously used for protein design or methods that performed well in the CAID2 competition.

In the case of a single disordered trypsin digestion site (Figure 4a), PIRATE and several of the prediction methods from the CAID2 competition can resolve a peak in disorder probability at or near the R336 digestion site. However, in a case where there are several trypsin digestion sites in proximity (Figure 4b), PIRATE is the only method that could cleanly resolve all three peaks of disorder probability within 5 residues of the trypsin digestion sites.

Researchers have shown that point mutations associated with the misfolding of the human alanine-glyoxylate aminotransferase (AGT) alter the protein’s sensitivity to trypsin digestion (Coulter-Mackie & Lian, 2008). For some mutations, this increased sensitivity was mitigated by binding to chemical chaperones, thus indicating that the mutations increased disorder at the experimentally validated trypsin digestions sites 36R and 122R. Notably, many of these point mutations were at distal locations to the trypsin digestion sites in terms of the primary sequence (Figure 5a and 5b). Given this observation, we sought to determine whether our approach would be sensitive enough to predict the effects of these distal mutations.

**Figure 5.**
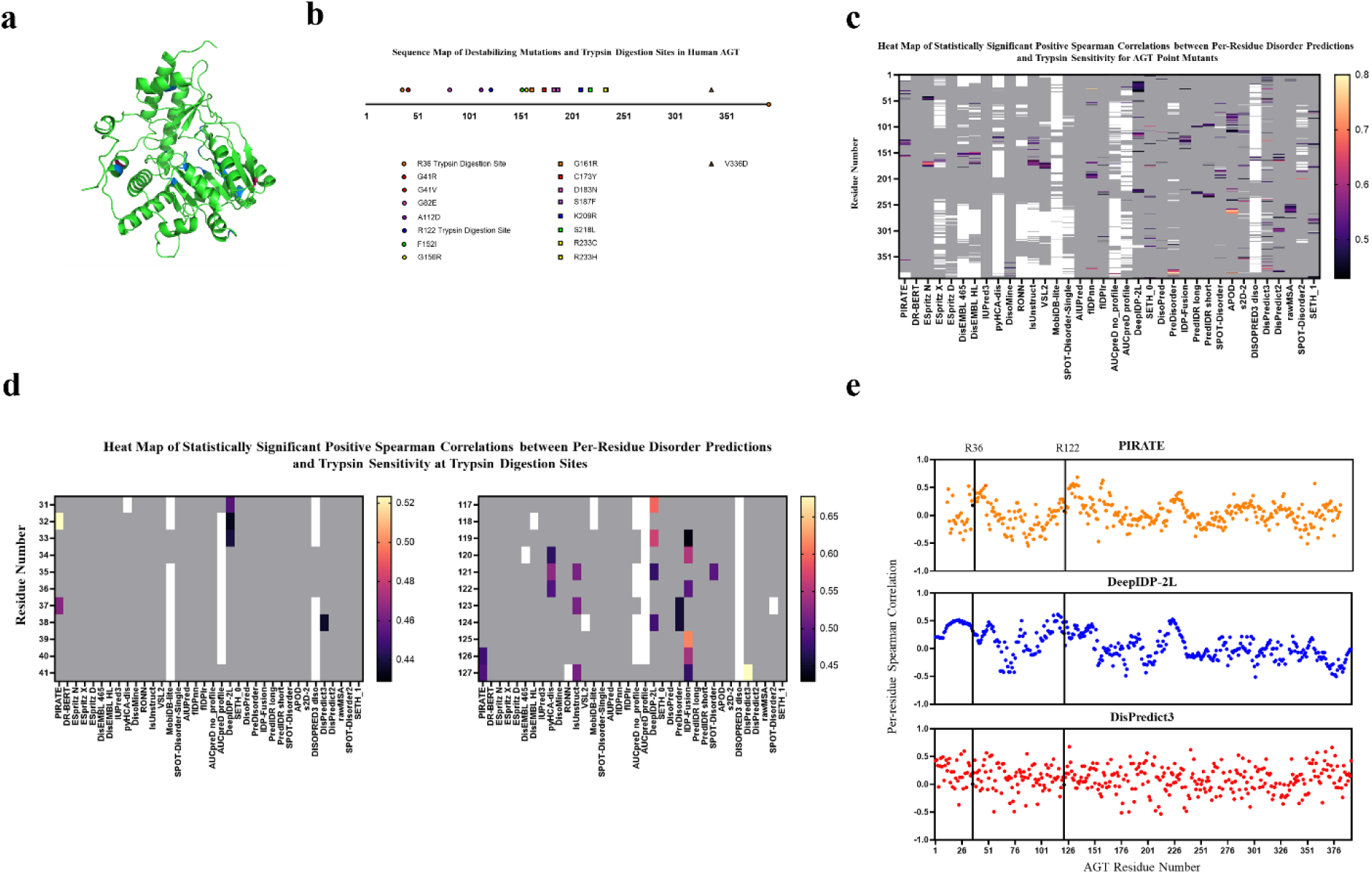
PIRATE Predictions Correlate with Trypsin Sensitivity in AGT Mutants. **(a)** An AF2 model for human AGT. Trypsin digestion sites are highlighted in red. Point mutation sites are highlighted in blue. **(b)** A sequence map of validated trypsin digestion sites and destabilizing AGT point mutations as described in Coulter-Mackie & Lian, 2008. **(c)** A heat map of statistically significant per-residue Spearman correlations between disorder predictions and trypsin sensitivity for AGT sequences. **(d)** A heat map showing statistically significant per-residue Spearman correlations between disorder predictions and trypsin sensitivity at the R36 and R122 trypsin digestion sites. **(e)** Plots showing the per-residue Spearman correlation between trypsin digestion scores and predicted disorder probabilities from PIRATE, DeepIDP-2L, and DisPredict3.

Through structural analysis of the WT protein, we rationalized how these mutations would impact structural stability. Glycine residues at positions 156 and 161 are essential for proper folding, as replacement with bulkier amino acids disrupts protein packing. Mutation of the glycine at residue 41 to either arginine or valine removes the interaction to its neighboring helix (exposing the trypsin digestion site R36). Mutation C173Y disrupts the protein’s tertiary structure as it causes internal steric clashes, while A112D introduces a charged residue into the hydrophobic core. R233 interacts with the N92 main chain but replacing it with histidine or cysteine causes misfolding due to shorter length of these side chains to maintain the interaction. Because the PIRATE methodology encodes sequence data in terms of local, regional, and global structural confidences as an intermediate step to making a disorder prediction, we expect it would predict the effect of these mutations on the order/disorder at the trypsin sites.

We predicted per-residue disorder probability for wild-type AGT as well as the fifteen point mutations illustrated in Figure 5b. For comparison, we also performed these predictions with DR-BERT and thirty-four other disorder prediction methods hosted on the CAID Prediction Portal. We looked for residue locations where there was a significant (p<0.05) positive Spearman correlation between predicted residue disorder and the trypsin digestion scores from Coulter-Mackie & Lian, 2008. Our hypothesis was that since these point mutations resulted in varying amounts of trypsin sensitivity, we should see a correlation between the per-residue disorder probability predictions and the experimental trypsin sensitivity at or near to the trypsin digestion sites.

When comparing the thirty-six methods of disorder prediction, we were surprised to find that many of the methods produced identical per-residue predictions for all the sequences at some locations. This results in a lack of ability to calculate a Spearman correlation at a residue and is illustrated as white on the heat map in Figures 5c and 5d. In the case of PIRATE, the white gaps on the sequence ends are due to PIRATE’s sliding window approach, which prevents disorder prediction for the 11 terminal residues. Residues where no significant correlation was found are labeled grey while statistically significant correlations are illustrated as shown on the color bar.

Only three of the disorder prediction methods evaluated showed significant positive correlation between predicted disorder and trypsin sensitivity within five residues of the trypsin digestion sites (Figure 5d). We plotted the per-residue Spearman correlations for these three methods (Figure 5e) and were pleased to see PIRATE produces peaks of Spearman correlation centered proximate to the locations of the trypsin digestion sites. The DeepIDP-2L (Tang et al., 2022) correlation plot also demonstrated distinct peaks of correlation, one of which is centered near to the R122 trypsin digestion site. DisPredict3 (Kabir & Hoque, 2024) showed non-localized correlation between disorder predictions and trypsin sensitivity, suggesting that it recognized increases in disorder caused by the destabilizing mutations, but that it would be ill-suited for identifying the specific protein regions associated with the misfolding phenotype.

### PIRATE Directed Evolution: Greedy Exploration simulation for automated protein design

To evaluate PIRATE’s capabilities for automated protein design, we developed a greedy exploration (GE) algorithm which we refer to as PIRATE directed evolution (PDE). In PDE, PIRATE is used iteratively to determine which residues to edit, and which mutations or deletions are most beneficial (Figure 6a). In its current implementation, PDE accepts only one modification to a single residue per cycle, selecting the change that results in the greatest mean disorder reduction across the entire protein. Mutations that produce a net increase in disorder either locally or globally are always excluded. The simulation terminates when the user-defined criteria are met, i.e., when the maximum number of mutations has been reached, or when the AI is unable to identify a beneficial mutation or deletion, causing it to self-terminate. The PDE interface offers several tunable parameters to guide the AI simulation protocol, including the specific sequence regions to modify (defined by residue number), the cumulative number of permitted modifications to the original sequence, the number of residues evaluated per iteration, and the number of disorder-reducing mutations to evaluate at each putative site per iteration.

**Figure 6.**
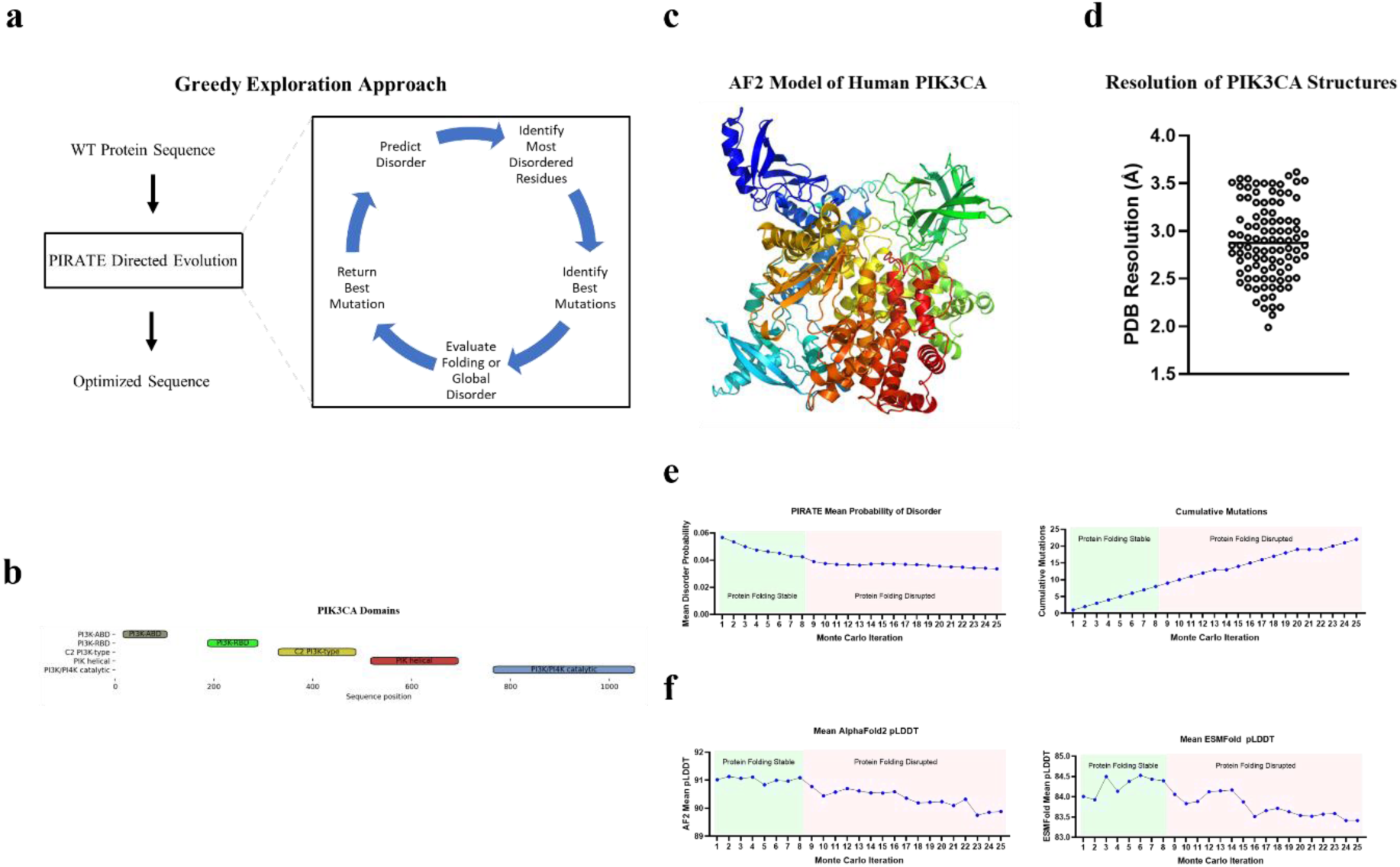
PIRATE Directed Evolution of PIK3CA. **(a)** Diagram of greedy exploration approach for reducing protein disorder. **(b)** Diagram of PIK3CA domains **(c)** AF2 protein model for PIK3CA **(d)** Average resolutions for PDB’s containing at least twenty percent of the full PIK3CA sequence **(e)** Plots showing the mean disorder and cumulative mutations across twenty-five directed evolution iterations. **(f)** Mean AF2 and ESMFold pLDDT at each iteration of the PIK3CA directed evolution.

We chose the human protein PI3KCA (Figure 6c) as a test case for evaluating PDE. PI3KCA is a high-value target where most attempts to experimentally determine protein structures with more than twenty percent of the canonical PI3KCA sequence have resulted in poorly resolved structures (Figure 6d). We allowed PDE to evaluate mutations at only 231 residues outside conserved domains (Figure 6b). During each iteration of the PDE simulation, PIRATE determined the five most disordered residues from the allowed mutations sites. For each of these residues, PIRATE screened all potential mutations, assessing their ability to reduce disorder. The five mutations that produced the greatest reduction in local disorder (averaged over the mutation site and the eleven adjacent residues on either side) were then evaluated for their ability to reduce the global protein disorder. From these five top-ranked mutations, PDE selected the one which produced the highest reduction in disorder across the entire protein.

The PDE was run for twenty-five iterations (Figure 6e) allowing the canonical PI3KCA sequence to accumulate twenty-two total mutations. Analyzing the sequences from each iteration, we found that both AlphaFold2 and ESMFold predicted that the mean pLDDT began to decrease after the eighth iteration (Figure 6f). This was an expected result as proteins require a certain amount of flexibility to accommodate folding.

We evaluated the mutations that occurred in the first eight iterations of the GE simulation by structural analysis. Structural analysis revealed several key mutations that we hypothesize are linked to the enhanced protein’s stability. Notable improvements included the addition of a putative hydrogen bond between H14E mutation and Q721 (Figure 7a), stabilization of secondary structure with P316Y (Figure 7b.), replacement of a polar threonine with the hydrophobic isoleucine in a hydrophobic pocket by the T322I mutation (Figure 7c), and the introduction of N107W which capped an otherwise rugged surface (Figure 7d).

**Figure 7.**
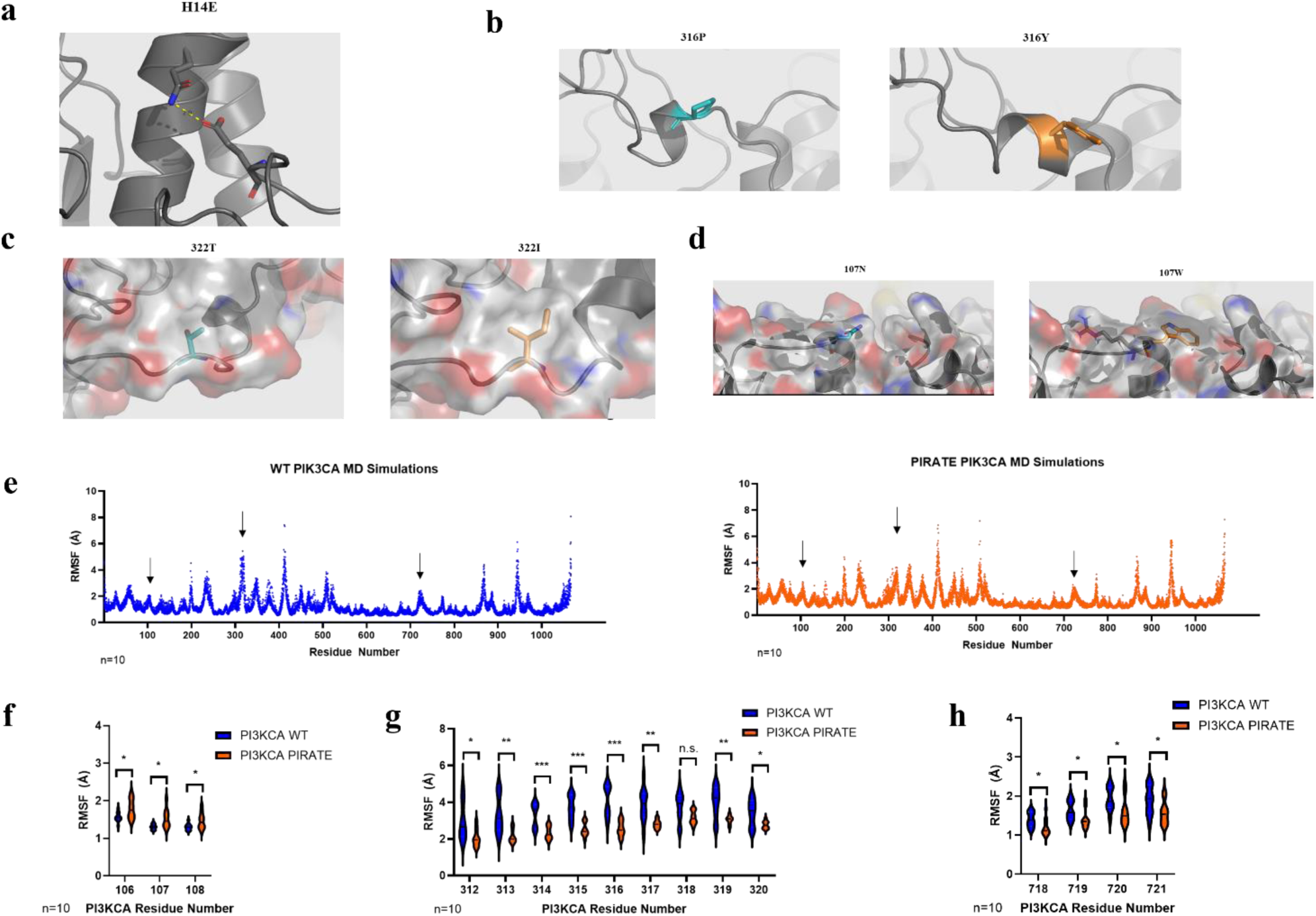
Analysis of PIRATE-Optimized PI3KCA. **(a)** H14E mutation introduces a hydrogen bond with Q721. **(b)** P316Y stabilizes and extends secondary structure. **(c)** T322I mutation replaces polar residue with hydrophobic residue inside hydrophobic pocket. **(d)** N107W mutation places a lid on a rugged protein surface **(e)** Molecular dynamics simulations show slightly increased movement at the site of N107W, stabilization of protein movement near mutation sites at residues 316 and 322, and in the region of the putative hydrogen bond formed by H14E. **(f)** Violin plot of RMSF values for MD simulations of WT and PIRATE-designed proteins at residues 106-108. **(g)** Violin plot of RMSF values for MD simulations of WT and PIRATE-designed proteins at residues 312-320. **(h)** Violin plot of RMSF values for MD simulations of WT and PIRATE-designed proteins at residues 718-721. Significance of unpaired, two-tailed student’s t-test is shown with one asterisk indicating p<0.05, two asterisks indicating p<0.01, and three asterisks indicating p<0.001.

To evaluate the stabilizing effects of these four mutations, we conducted ten replicate MD simulations for the WT protein and the variant containing the mutations. MD simulations showed reduced flexibility in the long loop spanning residues 312 to 322 as well as in the alpha helical region predicted to form a hydrogen bond with the H14E mutation (Figure 7e). The N107W mutation correlates with a small, but statistically significant increase in movement at residues 106-108 (Figure 7f). Structural analysis of N107W would suggest that the primary benefits of this mutation would be improved globularity and reduced probability of aggregation, so a small increase in movement is likely to be a beneficial tradeoff. Notably, we observed a large statistically significant reduction in C-alpha RMSF at all but one of residues 312-320 that we attribute to the stabilizing effect of the mutations at P316Y and T322I (Figure 7g). A similar reduction in C-alpha RMSF was detected at residues 718-721, which we attribute to the putative hydrogen bond between H14E and Q721 (Figure 7h).

## Discussion

In this study, we introduce PIRATE, a novel disorder prediction method specifically designed for practical tasks in protein engineering. PIRATE is based on three AF2 surrogate models that encode protein sequence as outputs of local, regional, and global confidence in structure. We show that the output from these surrogate models correlates intuitively with what we would expect for residues that are either modeled or unmodeled in published protein structures.

The approach was benchmarked using the most recently published CAID dataset. It performed slightly worse than the Alphafold2-RSA method, but slightly better than the published Alphafold2-pLDDT approach. By comparison to the best scoring methods in the CAID2 competition, PIRATE provides an excellent tradeoff between speed and performance, requiring only 1-2 seconds to make each inference on CPU with a Dell Latitude 5430 laptop (12th Gen Intel(R) Core (TM) i5-1235U, 32 GB Installed RAM).

Additionally, we demonstrate that PIRATE is highly capable at resolving the locations of small regions of disorder with biological significance such as cleavage sites. More importantly, we demonstrate that PIRATE possesses state-of-the art ability to predict how small perturbations to a sequence will impact structural disorder in distal locations. Further, we showed that PIRATE effectively elucidates the locations of trypsin digestion sites using only the identities of fifteen destabilizing point mutants and their corresponding trypsin sensitivity scores. Such an approach could potentially be used to probe the location and mechanism of any disorder-related biological process such as degron functionality or binding pockets. Inversely, we expect that a researcher could use PDE as a method for enhancing or reducing disorder-related biological function through distal point mutations.

Our results also demonstrate that many disorder prediction methods will make highly similar or identical predictions in the context of destabilizing point mutations. We feel that this underscores the advantages of using a method that encodes sequence data into local, regional, and global structural properties of the protein as an intermediate step before inference of per-residue disorder probability. Because our approach was trained across many AF2 predictions, it does a much better job inferring how a minor change at even a single location may alter disorder throughout the rest of the protein.

When we used PIRATE to iteratively make disorder-reducing mutations in non-conserved regions of PIK3CA, PIRATE rapidly found several mutations that structural analysis predicts will introduce a hydrogen bond, stabilize secondary structure, or smooth the surface of the protein. These improvements to protein rigidity were validated by improved stabilization of the corresponding regions during MD simulations.

While skilled structural biologists can replicate each of these protein engineering strategies, we are confident that it would take much more time for a human to design these stabilizing mutations compared to PIRATE. Additionally, it is less clear whether cooperative mutations such as P316Y and T322I would be human intelligible starting from a model of the WT protein. We anticipate that the usage of PIRATE as a foundational tool for engineering more stable proteins will enable us to overcome currently intractable problems in protein biochemistry since PIRATE can effectively identify strategies to stabilize disordered regions.

In light of these results, we would hypothesize that the current focus in disorder prediction may have become overly focused on achieving marginal improvements in binary disorder classification tasks with datasets created from measurements of proteins in non-biological contexts. Instead, we propose a shift towards training and benchmarking these methods to predict how minor changes to the protein sequence affect relative disorder across the entire protein. Additionally, assessing disorder prediction in more biologically relevant contexts may be key to unlocking the vast potential for protein design inherent to disorder prediction.

## Materials and Methods

### AlphaFold2 Datasets

AlphaFold2 predicted alignment error and pLDDT values were accessed from the AlphaFold Protein Structure Database (https://alphafold.ebi.ac.uk/) between October 2022 and January 2023. A list of UniProt accessions for the full dataset is available in Supplementary Data 1.

Because predicted alignment error is provided as an *N*x*N* array where *N* is the length of the target protein, we wanted to condense this into a single 1D array matching the length of the raw protein sequence. To do this, we took the mean of the predicted alignment array along each of the two axes and then took the average of these two mean values for each residue. These values were then scaled between 0.0 and 1.0 using the dataset min (1.24 Å) and max (30.06 Å) values. Residues were encoded using integers from 1 to 20 and then scaled using the dataset mean and std. deviation values such that the dataset mean was set to 0 and a standard deviation had a value of 1.0.

For creating the datasets to generate AFSM1/2, the sequences and their corresponding targets were randomly shuffled and then concatenated into 1D arrays. These were reshaped to an array with 4096 columns to accommodate an input shape of (1, 4096) for the UNet3+ architectures.

For creating the dataset used to train AFSM3, we downloaded the SEQRES for the PDB IDs listed in Supplementary Data 2. These sequences were shuffled and then appended to a 1D array one at a time. After each SEQRES was added to the array, we randomly selected a protein from the dataset for AFSM1/2 models. The longest continuous sequence where pLDDT was < 50 (minimum length of 25 residues) was added to the array. Residues from SEQRES were given a label of 1 while residues from sequence with a pLDDT < 50 were given a label of 0. We continued adding these low pLDDT residues to the array until the total number of low pLDDT residues was equal or greater than the number of SEQRES residues in the array. Then we would add the next SEQRES sequence until all the SEQRES sequences had been added to the array. This array was reshaped with 2048 columns to accommodate an input shape for the UNet++ model of (1, 2048).

### UNet Models

TensorFlow 2.8 was used to create all AFSM models for this paper. For AFSM1/2, a one-dimensional UNet3+ architecture was employed. The encoder part of the model was composed of convolution blocks composed of a single 1D convolution layer (kernel size 3, padding “same”) followed by a batch normalization (1D) layer and a ReLu activation layer. Down-sampling was performed by MaxPooling at each layer of the encoding network. The block used in each layer of the decoder network was composed of a Conv1DTranspose layer (stride 2, padding = “same”), a batch normalization layer, and a ReLu activation layer. As described in Huang et al., 2020, the skip full-scale skip connections were concatenated between respective layers using either MaxPooling or UpSampling followed by a convolution layer, a batch normalization layer, and a ReLu activation layer. The gross architecture of the network was five layers or blocks deep, the model input layer had a width of 128 units, and the output layer was a linear activation layer. We trained the AFSM1/2 models using a batch size of 8, learning rate of 0.005 with an Adam optimizer and early stopping with a patience of 3. Mean squared error was used as the loss metric.

The AFSM3 model architecture uses the same convolution and transconvolution blocks previously used for the encoding and decoding pathways in AFSM1/2. The model employs the dense nested connection structure described in Zhou et al., 2018 and as illustrated in Figure 1e which was shown to improve gradient flow. The model architecture was seven layers deep, and the input layer has a width of 64 units. The output layer was a sigmoid activation layer. We trained the model with a batch size of 4 using a learning rate of 0.0002 with Adam optimizer. Early stopping was employed with a patience of 3. Binary cross entropy was used as the loss metric.

### PIRATE Construction

We selected a dataset of 9,604 PDB structures containing only a single chain and consisting of proteins from the species used to construct the AlphaFold2 training dataset for the AFSM1/2 models. The list of PDB ID’s can be found in Supplementary Data 3. PDB’s and their mappings as to whether they were modeled in the published PDB structure were downloaded from the RSCB API (Berman, H. M., 2000) between 15July2023 and 26July2023. Residues that were present in the modeled PDB structure were given a label of 1 and those that were missing from the PDB structure were labeled as 0.

For each SEQRES, we encoded the structure properties of the model by inputting it into AlphaFold Surrogate Models and making a prediction of the normalized 1D predicted alignment error, the pLDDT, and the probability that a residue would be part of a SEQRES.

To input a sequence into the AlphaFold Surrogate Models, we initially encode the residues as integers. These are scaled (mean centered, standard deviation equal to 1) using the mean and standard deviation values of the corresponding training dataset. Since the input proteins are normally smaller than the input layer of the surrogate models, we pad the input with mirrored copies of the input sequence values on either side.

Each sequence was encoded using ASFM1/2/3 models and using ordinal values from 1-20. A sliding window of 23 consecutive positions is then passed over each of the encodings such that the value at position twelve represents the residue being evaluated for disorder with encodings from 11 residues on either side. The twenty-three values from each of the ASFM1/2/3 encodings and the ordinal encodings are then concatenated into an array of ninety-two values. These are the input features for a CatBoost model where the label (1 or 0) represents whether the residue represented by position twelve was modeled or unmodeled in the PDB structure.

In creating the dataset for training the CatBoost model, the initial protein sequences were shuffled. However, once the data were encoded and concatenated into ninety-two features for each residue, these data were handled sequentially to avoid data leakages.

Training data were down sampled using the AllKNN method from the imblearn library (Lemaître et al., 2017) to maximize the potential decision boundary across what we expected would be a very rugged landscape. The CatBoost model was tuned with the Optuna library (Akiba et al., *2019*) with an objective function maximizing Matthews Correlation Coefficient for the testing data. After 250 iterations of tuning, our best CatBoost classifier model had the following hyperparameters (iterations: 981, learning_rate: 0.09708238261031193, depth: 6, l2_leaf_reg: 0.000998506424925223, bootstrap_type: Bayesian, random_strength: 2.52125068317329e-08, bagging_temperature: 2.43641048326954, od_type: IncToDec, od_wait: 25). All other hyperparameters were left at their default settings.

For making disorder predictions, PIRATE encodes a raw protein sequence using the AFSM1/2/3 models and ordinal encodings. Ninety-two value representations are then created for each residue in the original protein sequence excluding the eleven residues on the N-term and C-term of the sequence since it is not possible. These are input into the CatBoost model for inference of the probability of class 0 (that a residue would be unmodeled in a PDB structure).

In Figure 2b, 2c, 2d, and Supplementary Figure 2, we show the mean predictions for each position in the twenty-three-value window for residues that were modeled or unmodeled in the structures listed in Supplementary Data 3. The sequence logos were made using the Python library logomaker (Tareen & Kinney, 2020).

### CAID2 Evaluation

PIRATE was evaluated based on per-residue predictions for the CAID2 dataset (Conte et al., 2023). For residues which could be classified, PIRATE achieved a performance of F1-max 0.8372, average precision score of 0.9059, and an AUC of 0.9333. In Figure 3, we compare these results to the results published in the CAID2 manuscript.

### DR-BERT Predictions

DR-BERT was downloaded from the project GitHub repository (https://github.com/maslov-group/DR-BERT) and used to make inferences locally. The model weights were accessed and downloaded as per their README instructions on 28May2024.

### Other Disorder Predictions

Other disorder predictions shown in the manuscript were generated by submitting sequences to the CAID prediction portal (https://caid.idpcentral.org/portal) between 12Aug2024 and 13Aug2024. TSV files resulting from these submissions are included in the project repository.

### AADC and EF1a Disorder Prediction

Canonical sequences for human AADC (UniProt ID: P20711) and human EF1a (UniProt ID: P68104) were evaluated by per-residue disorder probability in Figure 4.

### GraphPad Prism

GraphPad Prism 10 was used to prepare Figures 1b, 2b, 2c, 2d, 2e, 3, 4, 5b, 5c, 5d, 5e, 6d, 6e, 6f, 7e, 7f, 7g, and 7h. Prism was also used to generate the two-tailed Student’s t-test significances in Figure 7f, 7g, and 7h.

### Spearman Correlations

Spearman correlations reported in Figure 5 were generated using the SciPy (v1.11.1) library.

### AGT Disorder Prediction

Human AGT (UniProt ID: P21549) and the major allele point mutants characterized in Coulter-Mackie & Lian, 2008 were characterized for disorder using multiple disorder prediction methods. PIRATE and DR-BERT predictions were made locally, while the other methodologies shown in Figure 5 were generated using the CAID prediction portal. We encoded the trypsin sensitivities from Coulter-Mackie & Lian, 2008 using numbers to represent the data from Table 4 (no additions column) as that: - was scored as 0, -/+ was scored as 0.5, + was scored as 1, ++ was scored as 2, +++ was scored as 3, and ++++ was scored as 4. When we performed Spearman correlations between the trypsin sensitivity scores and the per-residue disorder predictions, we set 0.429 as the cutoff for statistical significance (p<0.05).

### PIRATE Directed Evolution

During the greedy exploration (GE) simulation to reduce disorder in human PI3KCA (UniProt ID: P42336), we allowed PIRATE to make point mutations to residues 1-15, 106-185, 290-328, 488-515, 695-763 in the canonical sequence. For each round of the GE simulation, PIRATE would identify the top 5 most disordered residues (among the residues it was permitted to mutate). At each of these five sites, the algorithm determined the best five mutations which would reduce predicted disorder the most at the residue site plus the eleven residues to either side of the mutation site (local disorder). The best five mutations from all five sites were then evaluated to determine which would reduce net disorder across the entire protein (global disorder) the most. Whichever mutation produced the largest net reduction in global disorder was selected for that iteration of the GE simulation. No mutations which produced increases in disorder locally or globally were permitted. In the case that a mutation satisfying this criterion could not be found, the simulation would self-terminate. We permitted the simulation to run for 25 consecutive iterations. The sequences and mutations from the PIRATE directed evolution are provided in Supplemental Data 5.

### AlphaFold2.3 models

All 3D models of the sequence variants proposed by PIRATE in this study were predicted by AlphaFold2.3 (https://github.com/google-deepmind/alphafold) (Jumper et al., 2021).

### MD Simulations

The WT PI3KCA protein and its variants were modeled using AlphaFold2, generating five structures for each. Only models with pLDDT scores exceeding 90 were considered and the structures with the lowest RMSD between WT and mutant were selected as the starting points for MD simulations. The C and N-termini were capped with an acetyl and a primary amide group, respectively, using PyMOL (Schroedinger, 2020). The protonation states of the amino acid residues were adjusted to reflect a pH of 7. The systems were prepared for MD simulations using the BioSimSpace library 2023.3.0 (Hedges et al., 2019), which in turn utilizes AmberTools20 (Case et al., 2020). The protein structures were parameterized using the AMBER ff14SB force field (Maier et al., 2015). Each system was placed in a cubic simulation box with a minimum distance of 12 Å between the protein and the box walls, and solvated with TIP3P water molecules (Jorgensen et al., 1983). Sodium (Na^+^) and chloride (Cl^-^) ions were added to neutralize the systems.

Subsequently, the systems underwent energy minimization, equilibration, and production MD simulations using AMBER 20.12.0 (Case et al., 2020). To ensure statistical robustness, ten independent simulations were performed for each protein. Initial velocities were assigned randomly, using a different seed for each replicate. In the energy minimization phase, solvent molecules were first relaxed while the protein coordinates were restrained with a force constant of 100 kcal mol^-1^ Å^-2^. This was followed by a full system minimization. Each minimization cycle consisted of 1000 steps of steepest descent, followed by 1000 steps of conjugate gradient minimization. Equilibration was carried out in three stages. Initially, the system was gradually heated from 0 K to 300 K over 250 ps with a 1 fs time step. Harmonic restraints were applied to the protein with a force constant of 100 kcal mol^-1^ Å^-2^. Afterwards, density equilibration was conducted in the NPT ensemble for 250 ps using a 2 fs time step, maintaining the same restraints. Finally, restraints were gradually released over three consecutive 250 ps NPT runs: initially restraining the protein backbone (10 kcal mol^-1^ Å^-2^), then only the C-alpha atoms (2 kcal mol^-1^ Å^-2^), and finally, releasing all restraints. This stepwise approach ensured stable equilibration over 1.25 ns. The production phase consisted of 100 ns of unrestrained NPT simulations at 300K with a 2 fs time step. Trajectories, energy values, and system coordinates were recorded every 20 ps.

Throughout equilibration and production, the temperature was controlled using the Langevin thermostat (Pastor et al., 1988) with a collision frequency of 5 ps^-1^. Pressure was maintained at 1 bar using the Monte Carlo barostat (Allen et al., 2017) with a pressure relaxation time of 1 ps. Periodic boundary conditions were applied, and long-range electrostatic interactions were computed using the particle mesh Ewald method (Darden et al., 1993). Van der Waals interactions were truncated at 10 Å and the pair list was updated at every simulation step. Bond lengths involving hydrogen atoms were constrained using the SHAKE algorithm (Ryckaert et al., 1977), with a tolerance of 10^-5^ Å. Additionally, rotational and translational motions of the system’s center of mass were removed every 1000 steps to prevent drift.

The analysis of the MD simulations was performed using MDAnalysis 2.5.0 (Michaud-Agrawal et al., 2011; Gowers et al., 2019). Prior to the analysis, the snapshots of the individual trajectories were aligned to the respective starting structures based on the C-alpha atoms. Conformational stability of the WT and mutant PI3KCA proteins was evaluated by monitoring the RMSF of the C-alpha atoms for all residues.

### PyMOL

PyMOL (Schrödinger & DeLano, 2020) was used to prepare Figures 5a, 6c, 7a, 7b, 7c, 7d.

### Code Availability

The code needed to reproduce key results from the manuscript as well as the core functionality for PIRATE can be accessed at https://github.com/Evotec-isRD/PIRATE.

## Acknowledgements

We would like to thank Sam Griffiths (Evotec Abingdon) for suggesting PI3KCA as a target for evaluating PDE in this study. We would also thank Mark Brooks (Evotec Princeton) for inspiring the creation of what ultimately became AFSM3.

---

**Supplementary Figure 1.**
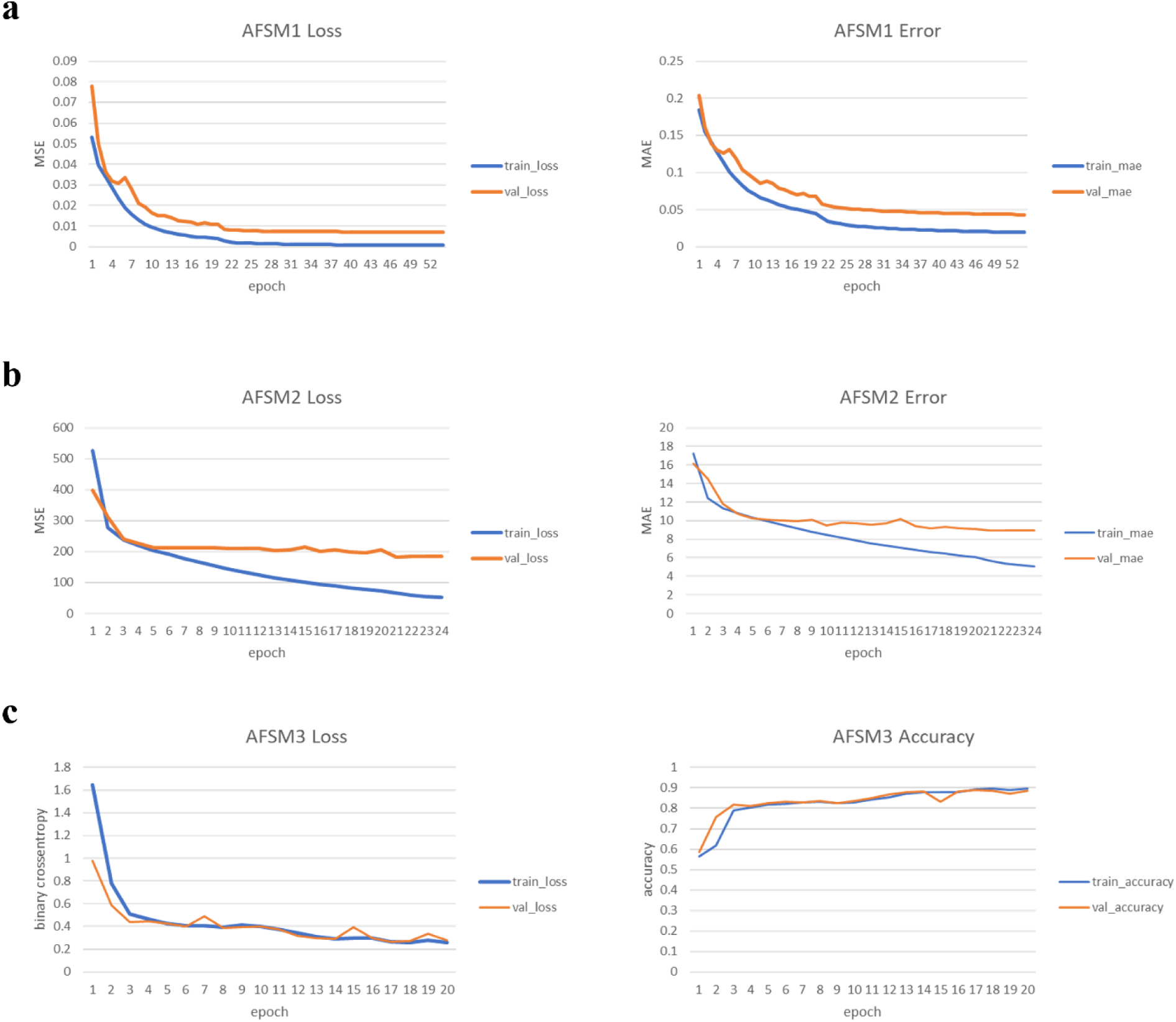
Training Curves for AlphaFold2 Surrogate Models. **(a)** Plots showing the loss (MSE) and mean absolute error curves generated while training AFSM1. **(b)** Plots showing the loss (MSE) and mean absolute error curves generated while training AFSM2. **(c)** Plots showing the loss (binary cross entropy), and accuracy curves generated while training AFSM3.

**Supplementary Figure 2.**
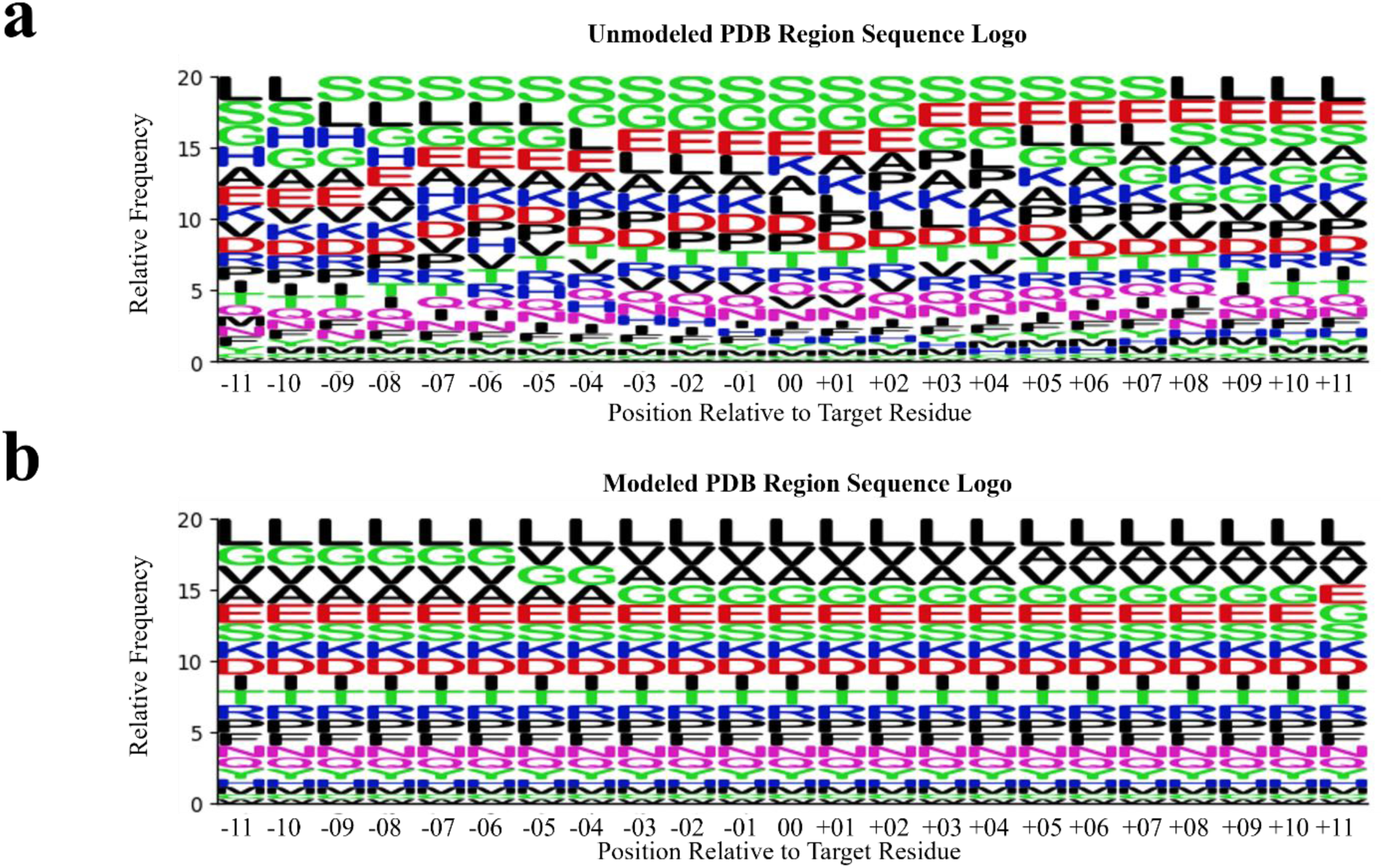
Sequence Logos for Modeled and Unmodeled Regions from PDB Dataset. **(a)** Sequence logo showing the mean representation for regions that were not modeled in the PDB dataset. **(b)** Sequence logo showing the mean representation for regions that were modeled in the PDB dataset.

## Notes

https://github.com/Evotec-isRD/PIRATE

### Competing Interest Statement

The authors have declared no competing interest.

### Summary of Updates

The figure conversion was redone to improve the readability of figures. Non-substantive edits were made to the manuscript text as well. ORCID's were added for three of the authors.

